# Simultaneous fMRI and eye gaze recordings during prolonged natural stimulation - a studyforrest extension

**DOI:** 10.1101/046581

**Authors:** Michael Hanke, Nico Adelhöfer, Daniel Kottke, Vittorio Iacovella, Ayan Sengupta, Falko R. Kaule, Roland Nigbur, Alexander Q. Waite, Florian J. Baumgartner, Jörg Stadler

## Abstract

Here we present an update of the *studyforrest* (http://studyforrest.org) dataset that complements the previously released functional magnetic resonance imaging (fMRI) data for natural language processing with a new two-hour 3Tesla fMRI acquisition while 15 of the original participants were shown an *audio-visual* version of the stimulus motion picture. We demonstrate with two validation analyses that these new data support modeling specific properties of the complex natural stimulus, as well as a substantial within-subject BOLD response congruency in brain areas related to the processing of auditory inputs, speech, and narrative when compared to the existing fMRI data for audio-only stimulation. In addition, we provide participants’ eye gaze location as recorded simultaneously with fMRI, and an additional sample of 15 control participants whose eye gaze trajectories for the entire movie were recorded in a lab setting — to enable studies on attentional processes and comparative investigations on the potential impact of the stimulation setting on these processes.

## Background & Summary

The investigation of the interplay of cognitive processes in a complex natural environment is an emerging research focus^1^. There is a growing number of publicly available datasets to facilitate this kind of research^2–5^ that primarily focus on aspects of processing sensory information in the visual domain. Complementing these resources, we previously released a high-resolution BOLD fMRI dataset on the processing of language and auditory information in a two-hour audio movie version of the Hollywood motion picture “Forrest Gump” — the *studyforrest* dataset (http://studyforrest.org)^6^.

One of the main challenges of working with natural stimulation paradigms is the presence of potential confounds that could impair the interpretability of results. Consider the case of an audio movie stimulus where the portrayal of emotional arousal happens to be correlated with loudness of the stimulus. In this case, it would be hard to dissociate the neural signatures of low-level stimulus properties from the neural representation of the perceived emotional state of a story character.

This problem can be addressed in two ways. First, by aiming to compose a complete description of the covariance structure of stimulus properties to allow for the discovery and quantification of potential confounds. Such a description should include low-level properties, often suitable for extraction with computer algorithms, and high-level features, such as a characterization of portrayed emotions^7^ that (still) requires human observers. Another approach is the combination of several stimuli that feature a different covariance structure of their particular properties in order to disentangle signatures of individual neural processes (see, for example, Hanke et al.^8^ for an extension of our original dataset with BOLD fMRI data on the perception of music for the same participants).

Here we present a further extension of the *studyforrest* dataset with data from 15 of the original participants. About a year after their initial participation, they *watched* and listened to the audio-visual “Forrest Gump” movie while fMRI data was being acquired, using the exact same movie cut and a stimulation that was synchronized with the previous acquisition. This additional stimulus delivers the same complex, two-hour story as the previous audio movie, but adds an enormous amount of visual information that detail and uniformly manifest new aspects of story content that were previously un(der)defined and left to a participant’s imagination — such as the specific composition of the visual scenery or the details of facial expressions.

This extension with an audio-visual movie enables the investigation of the representation of visual and multi-sensory input in a complex natural setting. Moreover, it facilitates comparative analyses with other existing datasets on movie perception^9^, as it is more similar in terms of its acquisition parameters and the nature of the stimulus content. At the same time, the combined dataset offers the unique opportunity to study the within-subject similarity of information representation across stimulus domains in the context of a complex and prolonged story narrative.

Attentional processes are likely to play a comparatively larger role in the selective processing of audio-visual vs. audio-only stimulation. With the goal of investigating the representation of information extracted from complex multi-sensory input, it is therefore of paramount importance to assess what particular aspects of the stimulus a participant is paying attention to at any point in time. In order to capture the location of the focus of attention, we recorded participants eye gaze coordinates for the entire duration of the movie simultaneously with the fMRI acquisition. This additional data modality enables further areas of investigation, such as which stimulus features drive attentional selection processes, but it also makes it possible to attempt to directly model aspects of the BOLD signal based on a behavioral measurement of visual spatial attention^10^. In order to be able to assess a potential influence of the stimulation setting in the scanner (small screen, participants are on their backs looking up), we include an additional sample of 15 participants who watched the movie in a lab setting on a larger screen while sitting in a comfortable chair.

In summary, this extension of the *studyforrest* dataset, together with its companion publication^11^, substantially increases the bandwidth of questions that can be investigated. Furthermore, the released data can be used to further improve the description of the stimulus itself: What object categories are being attended? When are eye movements made to track moving objects rather than switching the focus to a different object? Do participants at any given time explore the scenery, tune in on the dialog, or focus on a particular action being performed? Such future additions will help to further expand the scope and improve the interpretability of results.

## Methods

The information presented here is limited to a detailed description of aspects relevant to the simultaneous BOLD fMRI and eye gaze recording. For details on the general methods and a sample description, we refer the reader to the companion article^11^.

### Participants

All 15 participants described in Sengupta et al.^11^ also volunteered for the simultaneous fMRI and eye gaze recording in this study. Consequently, additional BOLD fMRI data for all participants have been made available previously^6,8^ and participant IDs are matched across all studies. One participant still had not seen the audio-visual “Forrest Gump” movie despite having participated in the audio-only movie study. The native language of all participants was German.

Although some participants posed substantial challenges for the eye tracking setup, such as partially obscured pupils due to long eye lashes or low contrast between iris and pupil due to insufficient lighting as a result of the head position in the head coil, no participant was excluded from the study. For those participants, the eye gaze recording was performed on a best-effort basis in order to maintain homogeneous procedures across participants while recording BOLD fMRI. Table 2 lists the affected participants. Participants in the fMRI experiment underwent a dedicated ophthalmologic exam to confirm normal vision^11^.

15 additional participants (age 19–30, mean 22.4, 10 females) volunteered for a separate eye tracking-only experiment. Two of those participants had never seen the movie “Forrest Gump” before. All participants had normal or corrected-to-normal vision.

Participants were fully instructed about the nature of the study, gave their informed consent for participation in the study as well as for public sharing of all obtained data in anonymized form, and received monetary compensation. This study was approved by the Ethics Committee of Otto-von-Guericke University (approval reference 37/13).

### Procedures

The procedures were highly similar to those described in Hanke et al.^6^. Before the scan, participants filled out a questionnaire on their basic demographic information and familiarity with the “Forrest Gump” movie.

fMRI data acquisition was split into two sessions that immediately followed each other on the same day. All eight movie segments were presented individually in chronological order with four segments in each session. Between sessions, participants left the scanner for a break with a flexible duration. On average, participants chose to continue with the second session after approximately 10 min.

At the start of the movie recording session, while auxiliary scans were performed, participants listened to music from the movie’s closing credits. During this time, the optimal stimulus volume was determined for each participant individually. Participants were instructed to “maximize the volume of the stimulus without it becoming unpleasantly loud or causing acoustic distortions by overdriving the loudspeaker hardware”. After the start of the movie, participants still had the ability to change the volume. Any adjustments were made within the first few seconds of the first movie segment; the volume remained constant across all movie segments otherwise.

Immediately prior to each movie segment, the eye tracker setup was calibrated (see eye tracking setup) and immediately followed by an accuracy validation. Directly after each movie segment ended, another gaze accuracy validation was performed.

After every movie segment, when the scanner was stopped and the gaze accuracy validation was completed, participants were asked to rate their experience during the preceding segment (“How deeply did you get into the story?”) on a scale from 1 (not at all) to 4 (a lot). Participants responded by pressing a button on a four-button response board with their right hand. This rating was followed by a brief break, and the recording continued as soon as the participant requested to continue by pressing a button.

Participants were instructed to inhibit any physical movements apart from eye movements, as best they could, throughout the recording sessions. Other than that, participants were instructed to simply “enjoy the movie”.

Procedures for the behavioral eye tracking session were practically identical, with the exceptions that there was no break between the first and second half of the movie and there was no dedicated audio volume calibration session. Instead, participants could manually adjust the volume throughout the experiment if they desired.

### Stimulus

Participants watched and listened to the movie “Forrest Gump” (R. Zemeckis, Paramount Pictures, 1994, dubbed German soundtrack). The stimulus source was the commercially available high-resolution Blu-ray disk release of the movie from 2011 (EAN: 4010884250916). Temporal alignment of the movie with the previously used^6^ DVD release was manually verified by audio waveform matching in order to guarantee synchronous stimulus time series between studies. The video track of the Blu-ray was extracted, re-encoded as H.264 video (1280×720 at 25fps), and muxed with the DVD’s dubbed German soundtrack using the MLT Multimedia Framework^12^.

Analog to the previous study, the audio track was processed by a series of filters in order to improve the audibility of the stimulus in the noisy environment of the scanner during echo planar imaging (EPI). Major noise components were located at 690 Hz and 1285 Hz as well as their harmonics at 2070Hz and 4140 Hz. More detailed information on the noise signature of the employed EPI sequence can be found in Angenstein, et al. (Figure 1)^13^.

**Figure 1.**
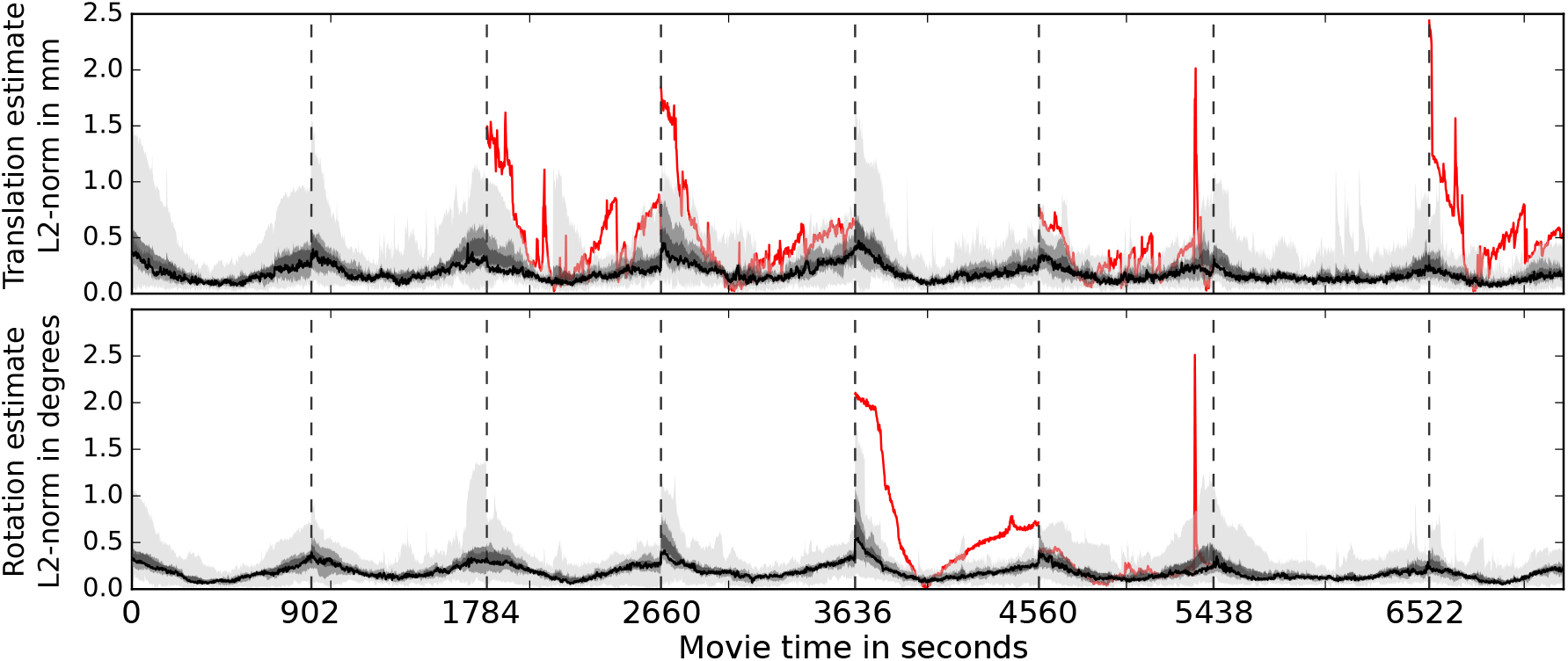
Summary statistics for head movement estimates across movie segments and participants. These estimates indicate relative motion with respect to template brain volume computed for each participants across all scans. The area shaded in light-gray depicts the range across participants, the medium-gray area indicates the 50% percentile around the mean, and the dark-gray area shows ± one standard error of the mean. The black line indicates the median estimate. Dashed vertical lines indicate run boundaries where participants had a brief break. The red lines indicate motion estimate time series of outlier participants. An outlier was defined as a participant whose motion estimate exceeded a distance of three standard deviations from the mean across participants for at least one fMRI volume in a run and exceeded a minimum translation of 1.5 mm or a rotation of 1.5°. For a breakdown of detected outliers, see Table 2.

For filtering, first a multi-band equalizer was used to implement a high-pass filter (-70 db attenuation in the 50 Hz band and −10 db at 100 Hz) to remove low frequencies that would have caused acoustic distortions in the headphones at high volume. This filter was followed by a dynamic range compressor (attack 128 ms, release 502 ms, 2:1 compression ratio above −18 db), a hard limiter (attack 128 ms, release 502 ms, cut off −12 db), and lastly a 3db volume gain. This chain implemented a conceptually similar filter as in the audio-only movie study^6^; however, the settings were optimized to the particular stimulation setup and environment. As a result, the filtered audio track used previously^6^ differs slightly from the audio-only movie stimulus, even in segments where the original soundtrack is completely identical between the audio-only and audio-visual movie.

Subsequently, the movie stimulus was shortened and cut into the same eight segments, approximately 15 min long each, as in the audio-only movie study. Additionally, the same fade-in/fade-out ramps were applied to the audio and video track (see Table 1 and Figure 3A in^6^).

**Table 1.** `participants.tsv` column description: Participant demographics and stimulus exposure. The original questions on the questionnaire were in German. To avoid clutter, only the translations are provided.

| Item | Description |
| --- | --- |
| participant_id | participant ID |
| gender | male: m; female: f |
| age | age group; five years width |
| handedness | right: r; left: l |
| hearing_problems_past | Have you had any hearing problems in the past (middle ear infections, otosclerosis, otospongiosis, hard of hearing, etc)? yes/no |
| hearing_problems_current | Do you have a current hearing deficit? yes/no |
| vision_problems_past | Have you had any vision problems in the past (strabismus, etc)? yes/no |
| vision_problems_current | Do you have a current vision deficit? yes/no |
| forrest_seen | Have you seen the movie “Forrest Gump” before? yes/no |
| forrest_seen_dist | How long since you last saw this movie? time coded in months. |
| forrest_seen_count | How many times have you seen this movie before? |
| forrest_seen_languages | What languages have you seen this movie in? |
| forrest_ad_known | Have you heard the audio movie version before? yes/no |
| forrest_av_rating | How did you like the movie?<br>bad: 1; rather bad: 2; rather good: 3; good: 4 |
| forrest_av_storydepth | How far did you delve into the story?<br>not at all: 1; a little: 2; rather deep: 3; very deep: 4 |
| forrest_av_fatigue | Did you get tired during the movie?<br>not at all: 1; a little: 2; rather tired: 3; very tired: 4 |
| forrest_av_feeling | How did you feel during the recording session?<br>bad: 1; rather bad: 2; rather good: 3; good: 4 |
| forrest_av_artist_count | In your opinion, music from how many artists was played in the movie? |

### Movie segment stimulus creation

As before, all video/audio editing was performed using the “melt” command-line video editor^12^ on a PC running the (Neuro)Debian^14^ operating system. As the actual commands are highly similar to those previously reported^6^, only key differences are described. The full source code for the stimulus generation is included in the data release (`code/stimulus/movie`).

All video files were created using a high-definition rendering profile (`-profile hdv_720 25p`). The watermark plug-in was used to replace the original black horizontal bars at the top and bottom of the movie frames with medium-gray bars of the same size in order to increase background illumination for a more pleasant experience (`-attach-track watermark:gray_bars_hdv720.png`).

Audio processing, as described above, was implemented using the following filter settings and utilizing two LADSPA-plugins (1197, Steve Harris, Ushodaya Enterprises Limited; 2152, Tom Szilagyi, Meltytech, LLC).

`# Multiband EQ`

`-attach-track ladspa.1197 0=-70 1=-10`

`# Compressor`

`-attach-track ladspa.2152 0=128 1=502 2=0 3=20 6=3`

`# Limiter`

`-attach-track ladspa.2152 0=128 1=502 2=0 3=-20 6=10`

`# Volume gain`

`-attach-track volume gain=3`

Video output specification options were:

`f=matroska acodec=libmp3lame ab=256k vcodec=libx264 b=5000k`

### Stimulation and eye tracking setup for fMRI acquisition

The visual stimulation setup was as described in the companion article^11^. Importantly, the movie was shown at a viewing distance of 63 cm in 720p resolution at full width on a 1280x1024 pixel screen that was 26.5 cm wide — corresponding to 23.75°x 13.5° of visual angle or 23.75° x 10.25° when considering only the movie content and excluding the horizontal gray bars.

Stimulation was implemented with PsychoPy v1.79 (with an early version of the MovieStim2 component later to be publicly released with PsychoPy v1.81)^15^ on the (Neuro)Debian operating system^14^.

Eye tracking was performed using monocular corneal reflection and pupil tracking with an Eyelink 1000 (software version 4.594) equipped with an MR-compatible telephoto lens and illumination kit (SR Research Ltd., Mississauga, Ontario, Canada). The temporal resolution of the eye gaze recording was 1000Hz. The eye tracking camera was mounted just outside the scanner bore, approximately centered, viewing the left eye of a participant at a distance of about 100 cm through a small gap between the top of the back projection screen and the scanner bore ceiling. An infrared light source, mounted slightly lower, illuminated the observed eye through the gap between the left side of the back projection screen and the scanner bore. The eye tracker was calibrated using a 13-point sequence that covered the entire display. For accuracy validation, participants had to fixate on the same 13 points, and offsets to the target coordinates were determined. Calibration was repeated as often as necessary until a sufficient consensus was achieved (see Technical Validation for accuracy estimates).

For participants 2, 10, and 20, the movie was presented vertically centered on the screen (the top movie pixel line below the gray horizontal bars was at y=239px). For all other participants, the movie display was shifted upwards by 171 px to yield a slightly more open eyelid and a better illumination of the pupil for improved eye tracking reliability.

Auditory stimulation was delivered through an MR confon mkII+ driving electrostatic headphones (HP-M01, MR confon GmbH, Magdeburg, Germany)^16^ fed from an Aureon 7.1 USB (Terratec) sound card through an optical connection. A participant’s head was fixed using a cushion with attached earmuffs containing the headphones. In addition, the participants wore earplugs. Headphones and earplugs each reduce the scanner noise by at least 20-30 dB, depending on the frequency.

### fMRI Response-/ synchronization setup

The TTL trigger signal emitted by the MRI scanner was fed to a Teensy3 microcontroller (`PJRC.COM`, LLC, Sherwood, OR, USA). The open collector signal from the response board (ResponseBox 1.2, Covilex, Magdeburg, Germany) was also fed into the Teensy3. A simple “teensydurino sketch” was used to convert the signals to USB keyboard events (“t”, “1”, “2”, “3”, “4”) at the stimulus computer.

Stimulus presentation and eye gaze recording were synchronized with the fMRI acquisition by automatically starting the eye gaze recording as soon as the stimulus computer received the first fMRI trigger signal. Moreover, the timing of subsequent trigger pulses was logged on the stimulus computer, and the onset of every movie frame (target frequency 25fps) was logged as part of the eye gaze recording via a custom log message to the eye tracker at the moment of the respective video buffer flip.

Due to the specific properties of the optical DVI extension system installed at the scanner, there was no direct synchronization of the video output of the stimulus computer with the refresh cycle of the projector. Moreover, it introduced a temporal offset and uncertainty between the recorded video update on the stimulus computer and in the eye tracker with respect to its actual appearance on the back-projection screen in the scanner. Separate measurements with a flickering test stimulus and a photo diode setup in the scanner-bore yielded a delay between target and actual onset time of 6-8 video refresh cycles (100-133 ms at 60Hz). No participant noted any audio-visual stimulus asynchrony.

### fMRI data acquisition

T2*-weighted echo-planar images (gradient-echo, 2 s repetition time (TR), 30 ms echo time, 90° flip angle, 1943 Hz/Px bandwidth, parallel acquisition with sensitivity encoding (SENSE) reduction factor 2) were acquired during stimulation using a whole-body 3Tesla Philips Achieva dStream MRI scanner equipped with a 32 channel head coil. 35 axial slices (thickness 3.0 mm) with 80x80 voxels (3.0x3.0 mm) of in-plane resolution, 240 mm field-of-view (FoV), anterior-to-posterior phase encoding direction) with a 10% inter-slice gap were recorded in ascending order — practically covering the whole brain. Philips’ “SmartExam” was used to automatically position slices in AC-PC orientation such that the topmost slice was located at the superior edge of the brain. This automatic slice positioning procedure was identical to the one used for scans reported in the companion article^11^ and yielded a congruent geometry across all paradigms.

The number of volumes acquired per movie segment was 451, 441, 438, 488, 462, 439, 542, and 338 volumes (for movie segments 1-8 respectively), and was therefore identical to the audio-only movie study^6^.

### Physiological recordings

Pulse oximetry and respiratory trace were recorded simultaneously with BOLD fMRI acquisition for the entire duration of the movie. The acquisition setup and the properties of the released data are described in the companion article^11^.

### Stimulation and eye tracking setup for in-lab acquisition

The stimulation setup for the in-lab acquisition was identical to the one at the MRI scanner except for the following differences: The screen for stimulus presentation was a BenQ XL2410T LCD monitor of size 522 x294 mm, displaying a resolution of 1920x 1080 px with a vertical refresh rate of 120Hz, and it was directly connected to the stimulus computer. The screen was positioned at a viewing distance of 85 cm, from which participants watched the movie while sitting in a slightly reclined chair — their head movements constrained by a U-shaped headrest that covered the back of the head. A different Eyelink 1000 with a standard desktop mount (software version 4.51; SR Research Ltd., Mississauga, Ontario, Canada) was used to record eye gaze coordinates at 1000 Hz using corneal reflection and pupil tracking of the left eye. In this configuration, the movie stimulus (excluding the gray horizontal bars at the top and bottom) subtended 34x15 “of visual angle.

### Code availability

All custom source code for data conversion from raw, vendor-specific formats into the de-identified released form is included in the data release (`code/rawdata_conversion`). fMRI data conversion from DICOM to NIfTI format was performed with heudiconv (https://github.com/nipy/heudiconv), and the de-identification of these images was implemented with mridefacer (https://github.com/hanke/mridefacer). The data release also contains the implementations of the stimulation paradigm.

## Data Records

This dataset is compliant with the Brain Imaging Data Structure (BIDS) specification^17^, which is a new standard to organize and describe neuroimaging and behavioral data in an intuitive and common manner. Extensive documentation of this standard is available at http://bids.neuroimaging.io. This section provides information about the released data, but limits its description to aspects that extends the BIDS specifications. For a general description of the dataset layout and file naming conventions, the reader is referred to the BIDS documentation. In summary, all files related to the movie data acquisition of a particular participant can be located in a `sub-<ID>/ses-movie/` directory, where ID is the numeric subject code.

All data records listed in this section are available on the OpenfMRI portal (dataset accession number: `ds000113d`) at http://openfmri.org/dataset/ds000113d as well as on Github/ZENODO^18^.

In order to de-identify data, information on center-specific study and subject codes have been removed using an automated procedure. All human participants were given sequential integer IDs. Furthermore, all BOLD images were “de-faced” by applying a mask image that zeroed out all voxels in the vicinity of the facial surface, teeth, and auricles. For each image modality, this mask was aligned and re-sliced separately. The resulting tailored mask images are provided as part of the data release to indicate which parts of the image were modified by the de-facing procedure (de-face masks carry a defacemask suffix to the base file name).

If available, complete device-specific parameter protocols are provided for each acquisition modality.

### Participant information

Data from the participants’ self-reports on basic demographic information and familiarity with the “Forrest Gump” movie are available in `participants.tsv`, a tabulator-separated value (TSV) file. This is an update of the demographics presented in the orginal data descriptor^6^. A description of all data columns is given in Table 1.

### Functional MRI

fMRI data files for the movie stimulation contain a `*ses-movie task-movie*_bold` pattern in their file name. Each image time series in NIfTI format is accompanied by a JSON sidecar file that contains a dump of the original DICOM metadata for the respective file. Additional standardized metadata is available in the task-specific JSON files defined by the BIDS standard.

### Physiological recordings

Time series of pleth pulse and respiratory trace are provided for all BOLD fMRI scans in a compressed three-column text file: volume acquisition trigger, pleth pulse, and respiratory trace (file name scheme: `_recording-cardresp_physio.tsv.gz`). The scanner’s built-in recording equipment does not log the volume acquisition trigger nor does it record a reliable marker of the acquisition start. Consequently, the trigger log has been reconstructed based on the temporal position of a scan’s end-marker, the number of volumes acquired, and with the assumption of an exactly identical acquisition time for all volumes. The time series have been truncated to start with the first trigger and end after the last volume has been acquired.

### Eye gaze recordings

Eye movement recordings are provided in two flavors: 1) raw, vendor-specific log files and 2) normalized gaze time series in a BIDS-compliant text file.

#### Raw eye gaze data in Eyelink ASCII format

For each movie segment, a dedicated log file with all raw data from the eye tracker is provided. It has been created by converting the original binary data file into ASCII text format using the vendor-supplied edf2asc tool and removing two header lines containing participant identifying information. Additionally, all files have been compressed with gzip. These files include annotated calibration parameters, acquisition time, gaze coordinates (x,y) in screen pixels, and pupil dilation; there are also integrated with basic information about automatically detected eye movements such as saccades, fixations, and blinks.

Files for eye gaze recordings performed simultaneously with fMRI data acquisition can be found at `sub-<ID>/ses-movie/func/sub-<ID>*_eyelinkraw.asc.gz`, whereas data from the separate lab experiment is available at `sub-<ID>/beh/sub-<ID>*_eyelinkraw.asc.gz`.

Note that all values in the data files from the in-scanner eye gaze recordings that specify information in units involving visual degrees cannot be used as such. Due to a misconfiguration, these measurements are based on a screen width of 37.6 cm instead of the actual 26.5 cm. Consequently, the actual value are about 70% smaller than the records in the files. This issue does not affect the actual eye gaze coordinates (recorded in screen pixels) nor does it affect the raw data files of the in-lab recordings.

#### Normalized eye gaze data

In addition, eye gaze coordinate time series are provided for all movie segments in a gzip-compressed tab-separated values text file. Each file contains four columns. The first two contain the *X* and *Y* coordinates of the eye gaze, followed by a pupil dilation measurement, and the numerical ID of the movie frame presented at the time of the measurement, as recorded by the eye tracker. The sampling rate is uniformly 1000 Hz, resulting in 1000 lines per second, with the first line corresponding to the onset of the movie stimulus.

These data have been normalized such that all gaze coordinates are in native movie frame pixels, with (0,0) being located at the top-left corner of the movie frame (excluding the gray bars) and the lower-right corner (again without the bar) located at (1280, 546) — correcting for the different display resolutions between in-scanner and in-lab recording and varying display location for the in-scanner recording. Moreover, the in-scanner recordings have been temporally normalized by shifting the time series by the minimal video onset asynchrony of 100 ms temporal normalization. The implementation of the normalization procedure can be found at `code/rawdata_conversion/convert_eyelink_to_cont`.

In-scanner eye gaze time series are located at `sub-<ID>/ses-movie/func/sub-<ID>*_recording-eyegaze_physio.tsv.gz` and in-lab recordings are at `sub-<ID>/beh/sub-<ID>*_recording-eyegaze_physio.tsv.gz`.

### Stimulus timing

Stimulus timing information for each recording segment are provided in BIDS’ `*_events.tsv` files. These six-column text files describe the `onset` and `duration` of each movie frame (`frameidx`) with respect to the MRI volume acquisition trigger signal (`lasttrigger`) as recorded on the stimulus computer (in contrast to the movie frame index column included in the eye tracking data files). Moreover, they also contain the timing log of the video `videotime` and audio `audiotime` stream of the stimulus movie, as reported by PsychoPy’s video component.

### Subjective story ratings

All participant ratings of subjective story depth are provided in a single JSON file for all movie segments combined: `ses-movie/sub-??_ses-movie_task-movie_bold.json` and for non-fMRI participants in `sub-??_task-movie_beh.json`.

## Technical Validation

All analyses presented in this section were performed on the released data in order to test for negative effects of de-identification or applied normalization procedures on subsequent analysis steps.

During data acquisition, technical problems were noted in a log. All known anomalies and their impact on the dataset are detailed in Table 2.

**Table 2.** Overview of known data acquisition anomalies (F: functional data, P: physiological recordings during fMRI session, ET: eye gaze recordings)

| Modality | Participant | Segment | Description |
| --- | --- | --- | --- |
| ET | 1 | 2 | > 30 % of no-signal samples |
| ET | 4 | 4 | > 30 % of no-signal samples |
| ET | 5 | 1-8 | > 30 % of no-signal samples |
| ET | 5 | 1,3,5 | initial average spatial error >73 px |
| ET | 5 | 3 | final average spatial error >93 px |
| ET | 10 | 5 | > 30 % of no-signal samples |
| ET | 10 | 1,6,8 | initial average spatial error >73 px |
| ET | 10 | 1,2,3,4 | final average spatial error >93 px |
| ET | 16 | 2 | > 30 % of no-signal samples |
| ET | 20 | 1-4,6-8 | > 30 % of no-signal samples |
| ET | 22 | 8 | initial average spatial error >27 px |
| ET | 23 | 2 | initial average spatial error >27 px |
| ET | 24 | 3,8 | initial average spatial error >27 px |
| ET | 29 | 7 | final average spatial error >81 px |
| ET | 30 | 8 | initial average spatial error >27 px |
| ET | 31 | 2 | initial average spatial error >27 px |
| ET | 36 | 2,3,6,7 | final average spatial error >81 px |
| P | 1 | 1-8 | technical failure, no data available |
| F | 5 | 1-8 | participant reported suboptimal sound quality |
| F | 9 | 5 | substantial motion >1.5 ° |
| F | 20 | 3,4,6,8 | substantial motion >1.5 mm |

### fMRI data quality

Participant movement is one of the key factors that negatively impact fMRI data quality. We estimated movement by aligning all time series images to a common participant-specific brain template image (rigid-body transformation implemented with FSL’s FLIRT). Figure 1 depicts separate summary statistics for estimates of translatory and rotary motion. With the exception of a few outliers (see Table 2), motion-related deviation from the reference image was found to be less than 1.5 mm and 1.5°.

For a description of the overall signal quality in terms of temporal signal-to-noise ratio, the reader is referred to the companion publication^11^.

Assessing fMRI data quality for natural stimulation paradigms beyond low-level metrics is a challenge due to the complexity of the employed stimulus. Here we employ two strategies to illustrate the presence of a rich signal in the presented dataset: BOLD responses correlated with a known sub-structure of the movie stimulus (portrayed emotions) and within-subject similarity of BOLD responses between this audio-visual movie stimulus and the previous audio-only movie study.

### BOLD response correlation with stimulus structure: portrayed emotions

One concern when using complex natural stimuli for research focusing on a particular cognitive function is the possibility that relevant neural signals are shadowed by other simultaneous cognitive processes. We tested whether it is possible to extract the signature of a particular aspect of brain activity under these conditions for the case of the response to emotional aspects of the stimulus.

In affective neuroscience, it is commonly assumed that observing emotional cues faces, words, characters, and social interactions elicits physiological brain responses similar to subjective experienced emotions — especially while viewing film clips^19^,^20^. Consequently, modeling BOLD responses in terms of the stimulus’ temporal dynamics with respect to portrayed emotions as well as their carrier modalities should reveal brain areas commonly associated with emotion processing. The pattern of activated brain areas need not be identical to those reported in studies using highly controlled experiments — as a more ecologically valid stimulus may be processed differently — but a substantial overlap is nevertheless to be expected.

Here, we used a previously published annotation of emotions portrayed in the movie stimulus^7^ to test for brain responses that occur over a variety of scenes, independent of character and context. To validate the selectivity of our procedure, we analyzed modality-specific emotion cues (visual or auditory). Additionally, we selected perception of self- versus other-referenced emotions in the movie as a higher-level aspect of affective processing. While the former analysis should yield modality-specific activations in sensory cortices, the latter analysis is expected to reveal areas implicated in social cognition — especially the representation of knowledge about others^21^.

In the present study, we used probabilistic indicators for the perception of emotion aspects based on the fraction of human observers reporting their presence for any time point in the movie (1Hz sampling rate)^7^. Each indicator encodes the inter-observer agreement in the interval [0, 1]: zero indicating no evidence for the presence of a property and 1 representing total agreement for the portrayal of a property across observers. The design matrix for a GLM time series model comprised of the following regressors: *high arousal, low arousal* (evidence for a deviation from an average or normal state of arousal), *positive valence, negative valence* (bipolar coding, analog to arousal), *self-directed emotion*, and *other-directed emotion*. In addition, the design matrix included regressors indicating evidence for a portrayed emotion from *verbal, non-speech audio* cue (separately), or via *facial expressions, gestures*, or other context cues. An additional boxcar regressor-of-no-interest indicated the presence of any speech in the stimulus, regardless of its emotional content. Lastly, a 6-parameter motion estimate was included.

A standard mass-univariate analysis was conducted with FEAT 6.0 from FSL 5.0.9^22^ that performed slice time correction, spatial smoothing (Gaussian FWHM 5mm), and highpass temporal filtering (Gaussian-weighted least-squares straight line fitting, with sigma 50s) as part of the pre-processing. Statistical analysis was carried out using FILM with local autocorrelation correction^23^. Time series analyses were performed for each movie segment separately, and were subsequently aggregated for each subject by averaging in a second-level analysis. Lastly, a group analysis was conducted testing for a mean effect in the group of all 15 subjects (random effects model; FLAME1) for the following contrasts: high>low arousal, positive>negative arousal, and self>other-directed emotion, as well as the respective reversed contrasts. Except for the negative>positive valence, there were significant clusters (threshold *Z* >2.3; *p*<0.05, corrected) of correlated signal for all conditions. The modulation of valence is rather inhomogeneous across movie segments, in particular for episodes with negative valence, and the approach of averaging results across segments might have impaired the analysis. Figure 2 shows the spatial distribution of these clusters for a subset of the results. Unthresholded maps for all contrasts are available on NeuroVault (http://neurovault.org/collections/1065).

**Figure 2.**
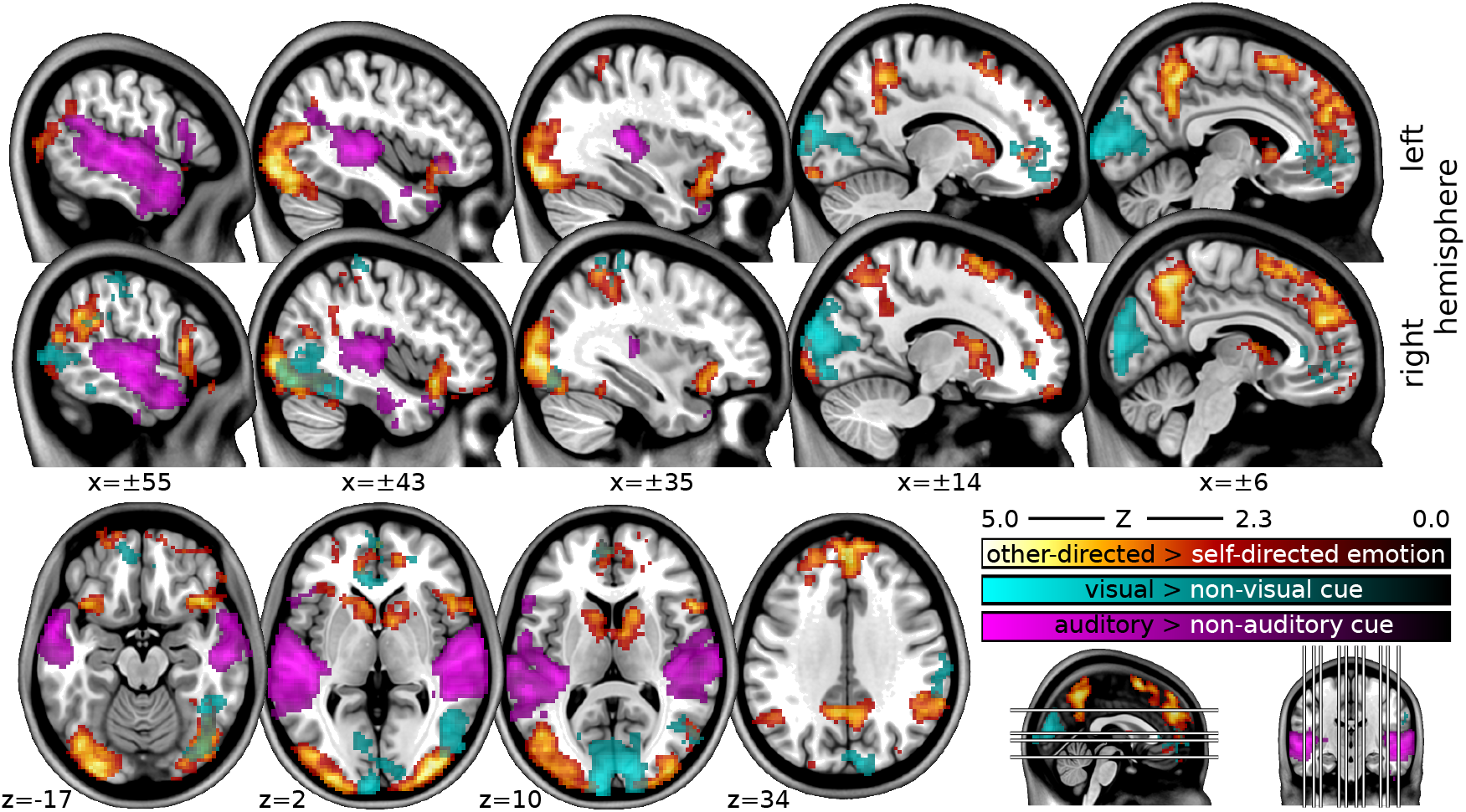
Mass-univariate regression group analysis of specific properties of portrayed emotion in the movie stimulus. Contrast 1: other-directed>self-directed emotion (red-yellow) lead to engagement of a large network including large parts of the dorsal medial cingulate gyrus, superior frontal gyrus, and precuneus, the latter being implicated in a variety of social cognition functions. Contrast 2-3: BOLD response correlations with portrayed emotions for auditory>non-auditory cues (purple) reveal primary auditory cortex and large portions of superior temporal gyri. In contrast, visual>non-visual cues (cyan) lead to pronounced activity patterns in occipital lobe comprising primary visual cortex and visual association areas. All coordinates are in MNI-space.

These results are notable for several reasons. First, group results for correlation with visual and auditory emotion cues show pronounced network activity in early visual areas as well as medial prefrontal areas (see Figure 2). The former is possibly an indication of a stimulus confound or a mediation of perceptual processes for emotional stimuli while the latter activation, comprising parts of anterior cingulate cortex as well as orbitofrontal (BA 24,32,10) regions, have previously been described by various studies as affective divisions of the medial frontal cortex^24–27^. Interestingly, auditory cue-related activity was completely isolated from these brain areas, leading to expected activation in auditory cortices.

Second, when contrasting the perception of other-directed emotions from self-directed emotions, activation patterns in the more dorsal parts of the cingulate cortex as well as areas in posterior parietal cortices appeared. The latter finding nicely supports meta-analytic findings regarding *theory-of-mind* (ToM) studies — suggesting that posterior parietal cortices are connected to the medial prefrontal cortex, thereby constituting a basic network for ToM^28,29^.

The selected results are evidence that these data contain reflections of emotional properties of the movie stimulus and that brain responses to individual aspects of portrayed emotions can be located by means of simple linear models. Furthermore, these results provoke a series of questions that could be studied using this dataset: Are dimensional or discrete emotion effects only visible in “emotion areas” or do they mediate perceptual processing? How does contextual information, such as social interactions or empathetic identification processes, influence the processing of perceived emotions? Additional relevant stimulus annotations are already available^7^ and further descriptions of this stimulus can enhance its utility as a reference for building upon previous movie stimulus studies on, for example: social perception^30^, facial expression^31^, fear^32^, humor^33^, sadness/amusement^34^, disgust/amusement/sexual arousal^35^, sadness^36^, happiness/sadness/disgust^37^, and emotional valence^38^.

### Within-subject response congruency to audio-only movie data

Except for the audio description content and low-level stimulus features due to equipment and filtering differences, the auditory aspect of the stimulation performed here was largely identical to the previous 7 Tesla fMRI acquisition^6^. Consequently, a substantial similarity of BOLD response time series between the two acquisitions is to be expected in brain areas associated with auditory and speech processing. We tested this with a voxelwise within-subject correlation analysis using the 14 participants for whom both 3Tesla and 7Tesla data were available. For each movie segment, the two respective time series images were motion-corrected (MCFLIRT) and spatially aligned to each other by means of a rigid-body transformation (FLIRT) and resliced to 2.5 mm isotropic resolution. Images were subsequently smoothed with a Gaussian low-pass filter (4mm FWHM; NiLearn). Voxel time-series were filtered by regressing out residual motion (implemented in PyMVPA using regressors estimated by the motion-correction algorithm) and band-pass filtered using a Butterworth filter (8^th^ order, −3 dB points at 16s and 250 s; implemented in SciPy). Following this movie-segment-wise pre-processing, voxelwise Spearmann rank-correlation coefficients were computed for corresponding movie segments from the audio-only and audio-visual experiment.

Correlation maps for all participants and movie segments were converted to standard Z-scores using Fisher transformation while normalizing to unit variance using 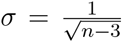, where *n* is the number of fMRI volumes in the respective movie segment. All Z-maps were then projected into MNI152 image space using participant-specific non-linear warps (FNIRT) and averaged across all participants and segments. Cluster-thresholding was performed using FSL’s *easythresh* tool, using the brain intersection mask of the 7Tesla (non-full brain) acquisition as the constraint.

Clusters of significant correlation between the two stimulation types could be observed in two large bilateral regions covering the auditory cortices, anterior and posterior STS, clusters located at Broca’s area (BA44/45, bilaterally; speech processing), and the precuneus, (all *Z* > 3.1, *p* < .05, cluster-corrected). At a more liberal threshold (*Z* > 2.3, *p* < .05, corrected), two large bilateral clusters also contain patches in the paracingulate gyrus/BA9 (implicated in story processing and mentalizing39). Notably, this also includes areas in the temporal occipital fusiform cortex — associated with face and scene perception — despite the lack of relevant visual stimulation in the audio-only movie experiment. Figure 3 shows the Z-map thresholded at *Z* > 2.3. The unthresholded map is available at http://neurovault.org/images/14268 for interactive inspection.

**Figure 3.**
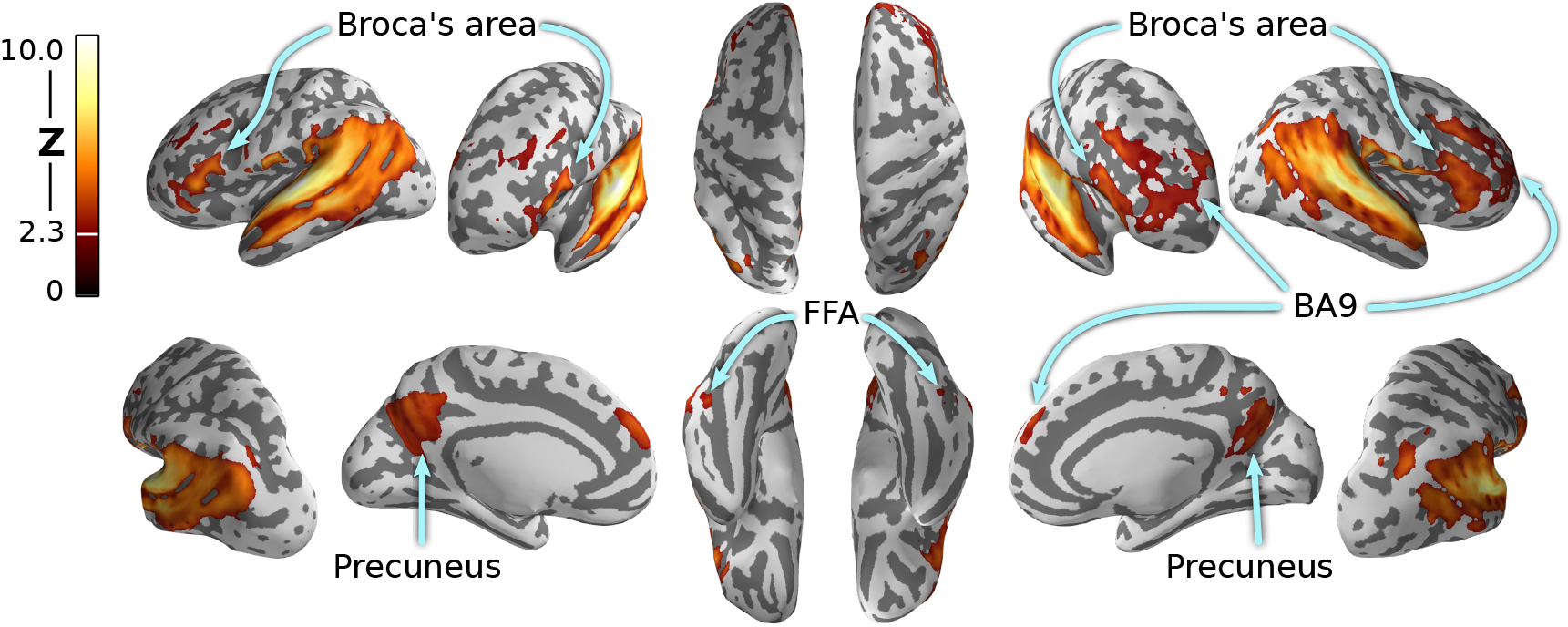
Average voxelwise within-subject correlation of BOLD time series between the previous audio-only movie experiment^6^ and the present audio-visual movie experiment. Correlations were Fisher-transformed into Z-score and averaged across movie segments (n=8) and participants (n=14). As expected, significant correlations are observed in brain areas associated with speech and story processing, but also in occipito-temporal cortex despite the lack of relevant visual stimulation in the audio-only experiment. Results are shown on the reconstructed surface of the MNI152 brain template.

### Eye tracking data quality

#### Timing precision

In order to assess the timing accuracy of frame presentation throughout the movie, for each movie segment, and for each participant, we extracted each movie frame’s onset time (as recored by the eye tracker from messages sent by the stimulus computer) and computed histograms of frame durations (Figure 4A). During the in-scanner experiment, the refresh rate of the presentation device was 60 Hz (1 refresh every ≈16 ms) and the movie frame rate was 25 Hz (1 frame every 40 ms). The overall distribution of durations peaked around two values (32 and 48 ms). More than 95% of the frames lasted for a duration of 32 ± 7 or 48 ± 7ms. During the laboratory experiment, the refresh rate of the presentation device was 120 Hz (1 refresh every ≈8ms). The overall distribution of durations peaked around two values (24 and 49 ms). More than 99% of the frames lasted for a duration of 24 ± 1 or 49 ± 1 ms. The overall timing stability of the in-lab sample was higher than the in-scanner acquisition. This is likely due to the lack of a direct synchronization signal between the stimulus computer and projector in the MRI-stimulation setup.

#### Signal loss

Due to the nature of the eye tracking technology, some signal loss is inevitable — primarily due to eye blinks; these lost samples are marked as `nan` (not-a-number) in the data files. For each participant, for each movie segment, the overall quantity of no-signal samples can be considered as an index of information quantity. For the majority of all participants, the signal loss is less than 10% (Figure 4B) across all movie segments. For 2 in-scanner participants out of the 15, more than 35% of the samples contain no signal (specifically: 85% and 36%). The amount of unusable samples for the remaining 13 participants ranges from less than 1% to about 15%. In-lab acquisitions yielded more contiguous data. Even the sessions with the highest amounts of loss included about 85% of usable data. Importantly, the movie content does not modulate the amount of no-signal samples: there is not a specific segment which shows, across the participants, significantly fewer usable samples with respect to the other segments. This holds for both the in-scanner and in-lab acquisitions.

**Figure 4.**
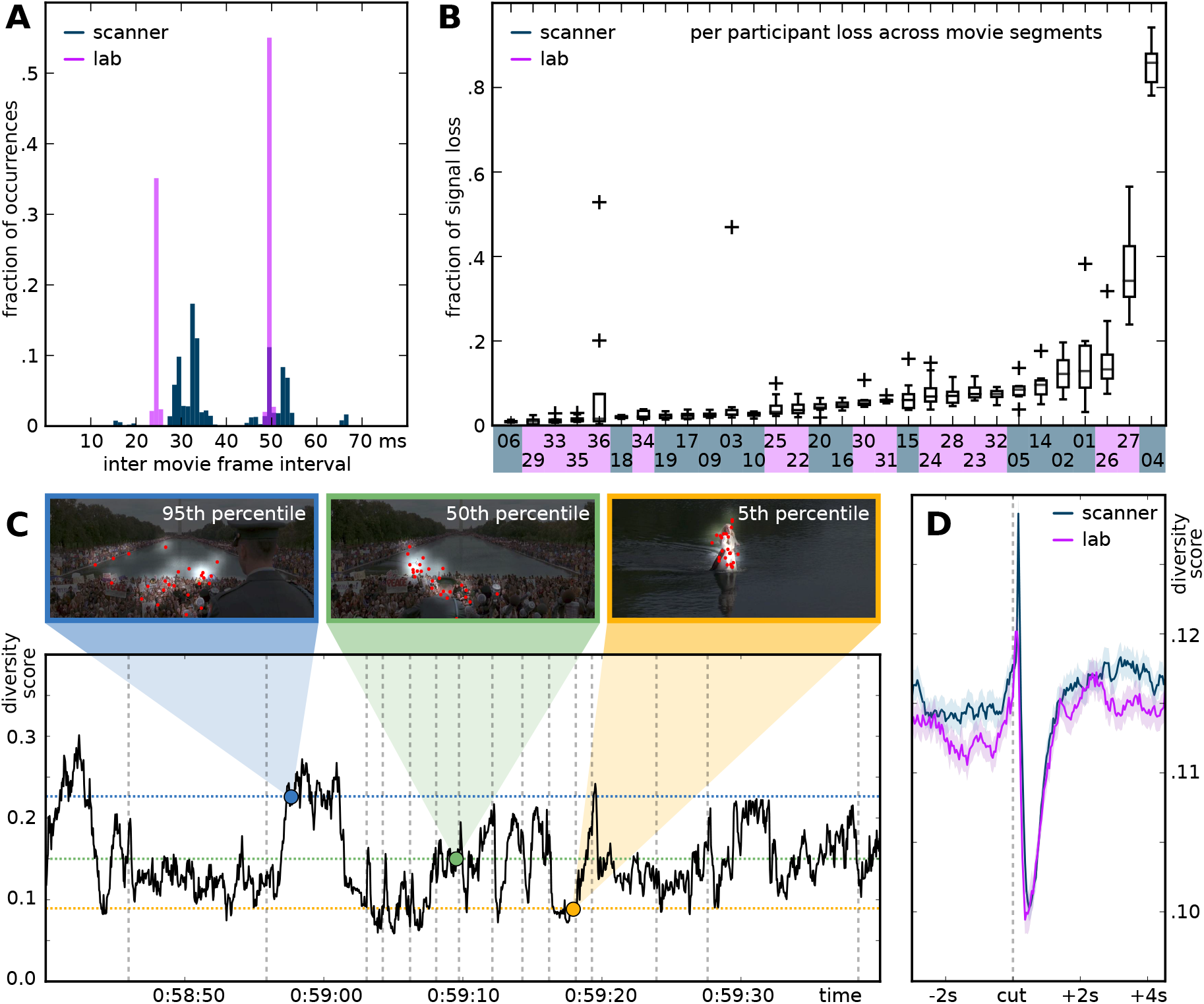
Technical validation of eye gaze recordings from the in-lab (purple curves) and the in-scanner (blue curves) sample. (A) Timing precision of movie frame presentation. The histogram shows the fraction of frame’s display durations across all participants and movie segments. (B) Fraction of signal loss per participants. The box plot show the distribution of signal loss across all movie segments, sorted by median loss. Sample association for individual participants is color-coded on the X-axis. (C) A one minute time series excerpt of the gaze diversity scores computed across all participants. The gray dotted vertical lines show the position of movie cuts. The horizontal blue line represents the 95th percentile for the full movie; green is for the 50th percentile, and the yellow for the 5th percentile. The three exemplary movie frames at the top show representatives of each chosen percentile containing the empirical gaze locations (red), and the brightness modulation on top of the movie frame content depicts the result of the Gaussian smoothing of the computed gaze distribution heat map. (D) The mean and standard error of all diversity score segments in temporal vicinity of a cut, separately computed for the in-scanner and in-lab participants (Y-axis conceptually similar, but not identical, to panel C). Participants 5, 10, 20, and 36 were excluded from the analyses presented in panels C and D due to low spatial accuracy of the eye gaze recordings (see Table 2).

#### Spatial accuracy

Spatial accuracy and reliability of eye gaze coordinates were estimated via dedicated 13-point test fixation sequences that were performed by each participant immediately prior to and after each movie segment. The unit of the presented results are movie pixels (reference movie frame size: 1280 x 546 px) to facilitate comparison between the in-scanner and in-lab recordings that employed different screen resolutions and physical dimensions.

##### In-scanner acquisition

Considering the average spatial error across any of the 13 calibration points on the screen prior to the start of the movie segments, the average spatial error across all participants and movie segments was 31 px (24/152/21 px median/max/std). The average error for measurement immediately following the movie segments increased to 41 px (36/170/26px median/max/std). The average within-subject increase of spatial error from the pre to the post movie segment test was 10 px (7/113/21 px median/max/std). This suggests an average spatial uncertainty of gaze coordinates of 40 px, corresponding to 0.75° of visual angle and a relatively stable coordinate uncertainty across the duration of the movie segments. Cases of extreme average error (more than two standard deviations from the mean) in the pre or post test are reported in Table 2.

##### In-lab acquisition

The analog estimates of spatial gaze coordinate error for the laboratory acquisition are generally lower and more homogeneous. The average pre-segment error is 17px (16/37/5px median/max/std), the post-segment error estimate is 37px (32/155/22px (median/max/std), and the average within-subject pre/post-segment error increase is 20px (14/143/24 px median/max/std). This suggests that more optimal optical conditions initially lead to more accurate gaze coordinates. However, accuracy degradation through the time course of the movie segments is more severe in comparison to the in-scanner acquisition — presumably due to superior head movement constraints in the head coil. Again, cases of extreme average error (more than two standard deviations from the mean) in the pre or post test are reported in Table 2.

#### Gaze distribution congruency across participants

In order to compare the in-lab and the in-scanner eye gaze recordings, we analyzed the spatial gaze distributions for every movie frame. Gaze distribution across all participants can be considered as an indicator of the magnitude of bottom-up attention modulation. Movie frames with clearly defined components of high saliency should lead to automatic attention capture and result in relatively synchronous eye movement across individuals. In contrast, movie frames without high-salience features, or temporally static movie frames after a period of initial exploration, should yield less synchronized and less coherent gaze locations across participants. We tested whether a quantification of between-subject gaze-diversity can be used to assess the synchronicity of our eye gaze recordings between the in-lab and in-scanner samples.

We computed a diversity score for every movie frame (duration 40ms) as the summed absolute difference of each cell of a uniform (flat) 2D gaze histogram and the empirical gaze histogram from the experiment (extent of a single histogram bin was 26×26 px). The latter was generated by composing a heat map from all gaze locations (one averaged location per participant per frame), smoothed by a Gaussian kernel with a standard deviation of 40px (the estimated spatial accuracy of the eye gaze recordings). High values indicate a high diversity of focus points across participants; low values indicate fewer and/or spatially adjacent focus points. Figure 4C illustrates this score with exemplary movie frames and their empirical gaze distribution for one average and two extreme cases from an arbitrarily selected one-minute segment of the stimulus movie.

Figure 4D depicts the temporal dynamics of this diversity score for 3 s prior and 4 s after each movie cut, averaged across the more than 850 cuts in the movie. A cut in a movie results in a sudden change of the visual stimulation and is assumed to yield maximum stimulus-driven attention modulation, leading to a decrease in the diversity score. We indeed observe such a global decrease with a local minimum at about 350ms after a cut. Moreover, Figure 4D documents highly similar temporal dynamics for the in-lab and the in-scanner sample, with the location of the respective minimum being only 50–80 ms apart. This difference could result from timing inaccuracies in the in-scanner sample, or from a slowed behavioral response in the scanner environment (participants resting on their backs). Prior the diversity minimum, we observe a diversity maximum at around 100 ms after a cut. One possible explanation is that some portion of the movie frames following a cut have multiple highly salient content locations to which saccades are performed immediately, but the saccade target differs across participants due to relatively equal salience. This question can be explored with a more detailed analysis of the provided eye gaze recordings.

Overall, we conclude that our diversity score captures relevant information about the gaze location congruency across participants, and that we observe highly similar temporal dynamics in both participant samples — indicating that these data are suitable for cross-sample comparisons.

## Usage Notes

The procedures we employed in this study resulted in a dataset that is highly suitable for automated processing. Data files are organized according to the BIDS standard^17^. Data are shared in documented standard formats, such as NIfTI or plain text files, to enable further processing in arbitrary analysis environments with no imposed dependencies on proprietary tools. Conversion from the original raw data formats is implemented in publicly accessible scripts; the type and version of employed file format conversion tools are documented. Moreover, all results presented in this section were produced by open source software on a computational cluster running the (Neuro)Debian operating system^14^. This computational environment is freely available to anyone, and it — in conjunction with our analysis scripts — offers a high level of transparency regarding all aspects of the analyses presented herein.

All data are made available under the terms of the Public Domain Dedication and License (PDDL; http://opendatacommons.org/licenses/pddl/1.0/). All source code is released under the terms of the MIT license (http://www.opensource.org/licenses/MIT). In short, this means that anybody is free to download and use this dataset for any purpose as well as to produce and re-share derived data artifacts. While not legally required, we hope that all users of the data will acknowledge the original authors by citing this publication and follow good scientific practise as laid out in the ODC Attribution/Share-Alike Community Norms (http://opendatacommons.org/norms/odc-by-sa/).

## Acknowledgments

We acknowledge the support of the Combinatorial NeuroImaging Core Facility at the Leibniz Institute for Neurobiology in Magdeburg. Michael Hanke was supported by funds from the German federal state of Saxony-Anhalt, Project: Center for Behavioral Brain Sciences. This research was, in part, co-funded by the German Federal Ministry of Education and Research (BMBF 01GQ1112, 01GQ1411) and the US National Science Foundation (NSF 1129855, 1429999) as part of two US-German collaborations in computational neuroscience (CRCNS).

MH conceived the study, acquired the data, performed and coordinated the validation analysis, and wrote the manuscript. NA acquired the in-lab eye tracking data and contributed to the manuscript. DK performed the inter-subject eye gaze congruency analysis. VI performed the quality control for the eye tracking data and contributed to the manuscript. AS contributed to the fMRI data acquisition. FRK performed quality control analysis for the fMRI data and contributed to the manuscript. RN contributed to the fMRI data emotion regression analysis and to the manuscript. AQW contributed to the in-lab eye tracking setup and to the manuscript. FB helped develop the initial in-scanner eye tracking setup. JS was the data acquisition lead and developed the in-scanner response synchronization setup.

## Competing financial interests

The authors declare no competing financial interests.

